# Multimodal normative modeling in Alzheimer’s Disease with introspective variational autoencoders

**DOI:** 10.1101/2024.12.12.628273

**Authors:** Sayantan Kumar, Peijie Qiu, Braden Yang, Abdalla Bani, Philip R.O Payne, Aristeidis Sotiras

## Abstract

Normative models in neuroimaging learn patterns of healthy brain distributions to identify deviations in disease subjects, such as those with Alzheimer’s Disease (AD). This study addresses two key limitations of variational autoencoder (VAE)-based normative models: (1) VAEs often struggle to accurately model healthy control distributions, resulting in high reconstruction errors and false positives, and (2) traditional multimodal aggregation methods, like Product-of-Experts (PoE) and Mixture-of-Experts (MoE), can produce uninformative latent representations. To overcome these challenges, we developed a multimodal introspective VAE that enhances normative modeling by achieving more precise representations of healthy anatomy in both the latent space and reconstructions. Additionally, we implemented a Mixture-of-Product-of-Experts (MOPOE) approach, leveraging the strengths of PoE and MoE to efficiently aggregate multimodal information and improve abnormality detection in the latent space. Using multimodal neuroimaging biomarkers from the Alzheimer’s Disease Neuroimaging Initiative (ADNI) dataset, our proposed multimodal introspective VAE demonstrated superior reconstruction of healthy controls and outperformed baseline methods in detecting outliers. Deviations calculated in the aggregated latent space effectively integrated complementary information from multiple modalities, leading to higher likelihood ratios. The model exhibited strong performance in Out-of-Distribution (OOD) detection, achieving clear separation between control and disease cohorts. Additionally, Z-score deviations in specific latent dimensions were mapped to feature-space abnormalities, enabling interpretable identification of brain regions associated with AD pathology.

## 1 Introduction

Brain disorders like Alzheimer’s Disease (AD) impact millions worldwide, significantly reducing the quality of life for patients and their families (Kumar et al., 2021; Richards and Brayne, 2010; Yang, Earnest, Kumar, Gordon, and Sotiras, 2024; Yang, Earnest, Kumar, Kothapalli, et al., 2024). While treatments like medication and therapy can alleviate symptoms, research often relies on case-control approaches that emphasize group averages, assuming uniform disease impacts across patients (Verdi et al., 2021). However, individual deviations from group means can provide critical insights for patient stratification and precision medicine (Marquand et al., 2016, 2019). Understanding heterogeneity within clinical cohorts at the individual level can enhance diagnosis and treatment strategies for AD (Jack et al., 2010; Kumar, Earnest, et al., 2024; Kumar, Oh, et al., 2024; Kumar, Yu, et al., 2024; Li et al., 2022). Normative modeling addresses this by capturing individual variability instead of group averages (Kia and Marquand, 2019; Marquand et al., 2019; Verdi et al., 2023). Traditional methods like Gaussian regression and w-scores often focus on univariate analysis, overlooking multivariate data interactions (Earnest et al., 2024; Lee et al., 2022; Loreto et al., 2024; Verdi et al., 2023). Recent deep learning frameworks, such as autoencoders and variational autoencoders (VAEs), address this limitation by modeling complex, nonlinear multivariate interactions (Kumar, Payne, and Sotiras, 2023; Kumar, Payne, and Sotiras, 2023; Lawry Aguila et al., 2023). VAEs encode input data into a lower-dimensional latent space to identify out-of-distribution (OOD) deviations and create subject-specific variability maps (Loreto et al., 2024; Verdi et al., 2023).

Despite their advantages, most VAE-based models are limited to single-modality data, missing the complementary insights provided by multimodal neuroimaging (Kumar, Earnest, et al., 2024; Kumar et al., n.d.; Lawry Aguila et al., 2022; Pinaya et al., 2019, 2021). This limitation is critical for multifactorial disorders like AD, where multiple interacting pathological processes contribute to progression. Multimodal integration of neuroimaging techniques such as MRI and PET is essential for capturing disease effects with varying sensitivities and advancing normative modeling frameworks.

Multimodal VAE-based deep normative models suffer from two major limitations. The core idea behind the normative approach is that the VAE is trained to reconstruct data from healthy controls, resulting in inaccurate reconstructions (higher reconstruction errors) when applied to AD patients. These errors can be used to measure each subject’s deviation from the norm and identify outliers. However, standard VAE models often struggle to accurately capture the distribution of healthy data, leading to large residuals and false positives for healthy controls. Prior research has shown that standard, variational, and adversarial autoencoders often fail in Out-Of-Distribution (OOD) detection tasks where normal and abnormal distributions overlap significantly (Bercea et al., 2023).

A further limitation of multimodal VAE-based normative models is their method of aggregating information across modalities, typically by estimating a product (Product-of-Experts, PoE) or an average (Mixture-of-Experts, MoE) of unimodal latent posteriors. This approach often leads to uninformative joint latent distributions, which hinders the accurate estimation of subject-level deviations. In PoE, a highly precise modality can dominate the joint inference, reducing the effectiveness of less precise modalities (Wu and Goodman, 2018). On the other hand, MoE distributes its density across all experts, resulting in a joint posterior no more refined than that of any single modality (Shi et al., 2019).

To address these limitations, our contributions are four-fold. First, we introduce a novel multimodal soft-introspective VAE (mmSIVAE) for normative modeling across multiple neuroimaging modalities. Soft-IntroVAE (SIVAE) (Daniel and Tamar, 2021) is a state-of-the-art framework for image generation and out-of-distribution (OOD) detection, training a VAE adversarially to enable the encoder to distinguish between generated and real data. We extend SIVAE to handle multiple modalities and provide a theoretical analysis of Nash equilibrium between the encoders and decoders. We hypothesize that our multimodal introspective VAE framework will enhance normative modeling by capturing a more precise representation of healthy anatomy, both in the latent space and in reconstructions. Second, we adopted the Mixture-of-Product-of-Experts (MOPOE) approach, combining the strengths of POE and MOE to aggregate multimodal information effectively in the latent space. Our main objective in this study to assess whether our proposed mmSIVAE with MOPOE aggregation can effectively learn the training distribution of a healthy population, accurately reconstruct inputs from their latent representations, and reliably detect outliers. Third, mmSIVAE quantified the deviation of AD patients based on the shared latent space learnt by MOPOE and label subjects with abnormal (statistically significant) deviations as outliers. We hypothesize that normative modeling with latent deviations better captures outliers compared to the baseline methods. We clinically validated the latent deviations to check if they were sensitive to the different AD stages and if they were significantly associated with cognition. Finally, clinical interpretation of deviations in the latent space is challenging. Towards this end, we identified latent dimensions with abnormal deviations, mapped them to feature-space deviations and analyzed regional brain deviations

## 2 Background

Let *X* = [*x*_1_, *x*_2_, …*x*_*N*_] be a set of conditionally independent N modalities. The backbone of our model is a multimodal variational autoencoder (mVAE), a generative model of the form 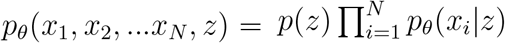, where z is the latent variable and *p*(*z*) is the prior. mVAE optimizes the ELBO (Evidence Lower Bound) which is a combination of modality-specific likelihood distributions *p*_*θ*_ (*x*_*i*_|*z*) and the KL divergence between the approximate joint posterior *q*(*z*|*X*) and prior *p*(*z*).

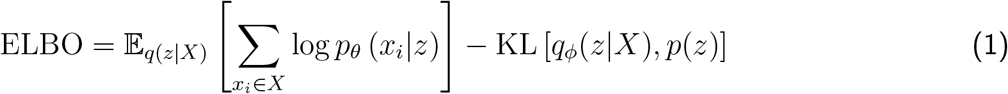

### 2.1 Soft-Introspective VAE (SIVAE)

#### Introspective VAE

Unlike the standard VAE, which optimizes a single lower bound, Introspective VAE (IntroVAE) (Huang et al., 2018) incorporates an adversarial learning strategy similar to that used in Generative Adversarial Networks (GANs). In this framework, the encoder’s objective is to maximize the KL divergence between the generated (fake) image and the latent variable while minimizing the KL divergence between the actual image and the latent variable. Simultaneously, the decoder seeks to challenge the encoder by minimizing the KL divergence between the generated image and the latent variable. The learning objectives for the Encoder and Decoder in IntroVAE are as follows:

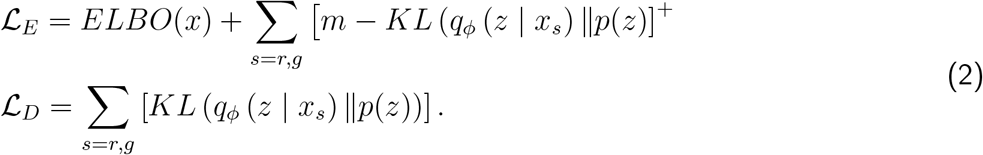

where *x*_*r*_ is the reconstructed image, *xg* is the generated image, and *m* is the hard threshold for constraining the KL divergence.

#### Soft-Introspective VAE

A key limitation of IntroVAE is its use of a hard threshold m to constrain the KL divergence term. Soft-Introspective VAE (SIVAE; Daniel and Tamar, 2021) argue that this design significantly reduces the model’s capacity and can lead to vanishing gradients. To address this issue, SIVAE proposes utilizing the entire Evidence Lower Bound (ELBO) rather than just the KL divergence, employing a soft exponential function in place of a hard threshold. The learning objective (i.e., loss function) for SIVAE is as follows:

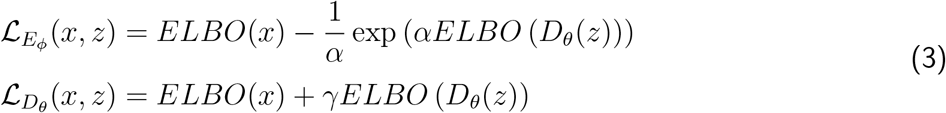

### 2.2 Approximating joint posterior in the latent space

#### Product-of-Experts (POE)

The approximate joint posterior *q*_*P oE*_(*z* | *x*) can be estimated as the Product of Experts (POE), where the experts are the unimodal approximate posteriors *q*_*ϕ*_(*z* | *x*_*i*_) and *p*(*z*) and *p*(*z*) is the “prior” expert.

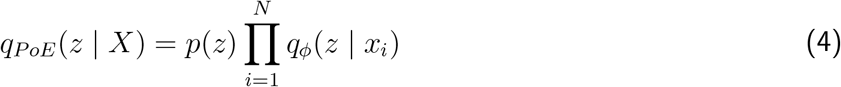

The product distribution required above are not in general solvable in closed form. However, if we approximate both *p*(*z*) and 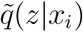 as Gaussian, a product of Gaussian experts is itself Gaussian with mean *µ* = (∑_*i*_ *µ*_*i*_ * *T*_*i*_)(∑_*i*_ *T*_*i*_)^−1^ and variance *σ* = (∑_*i*_ *T*_*i*_)^−1^ where *µ*_*i*_ and *σ*_*i*_ are parameters of the i-th Gaussian expert and 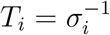.

The challenge of approximating inference distribution with POE is that if the inference distribution for a particular modality (expert) is very sharp, then the joint distribution will be dominated by it. In other words, the overconfident but mis-calibrated experts may bias the joint posterior distribution which is undesirable for learning informative latent representations between modalities.

#### Mixture-of-Experts (MOE)

Another way to approximate joint inference is to use the mixture of experts (MoE) form, where the approximate posterior of the joint is represented by the sum of the unimodal posterior distributions. Since MoE is the sum of each expert and not product, the joint posterior distribution is not dominated by experts with high precision as in PoE, but spreads its density over all individual experts. In the MoE setting, each uni-modal posterior *q*_*ϕ*_(*z* | *x*_*i*_) is evaluated with the generative model *p*_*θ*_(*X, z*) such that the ELBO becomes:

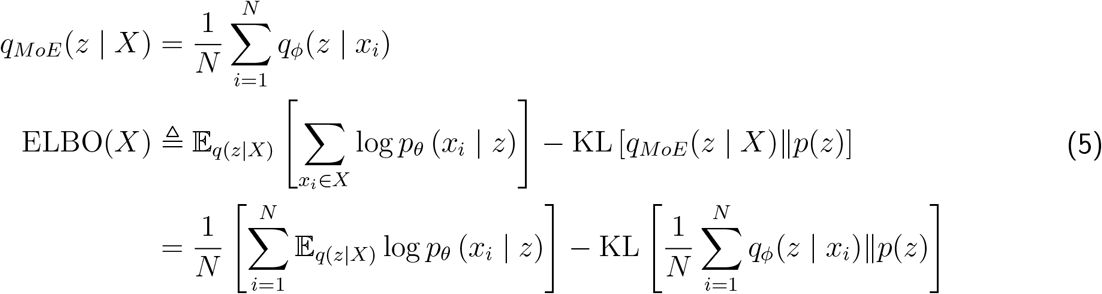

But this approach only takes each unimodal encoding distribution separately into account during training. The aggregation of experts in MoE does not result in a distribution that is sharper than the other experts. Therefore, even if we increase the number of experts, the shared representation does not become more informative as in PoE. Thus there is no explicit aggregation of information from multiple modalities in the latent representation for reconstruction by the decoder networks.

## 3 Proposed methodology

In this section, we introduce multimodal soft-introspective VAE (mmSIVAE), extending SIVAE (Daniel and Tamar, 2021) to the multimodal setting. First, we present the theoretical analysis of mmSIVAE, followed by modeling of the joint latent posterior using the Mixture-of-Product-of-Experts (MOPOE) technique. Next, we discuss how mmSIVAE can be adopted for multimodal normative modeling which includes model training, inference and calculation of subject-level deviations.

Following Equation (3), the encoder and decoder loss functions for mmSIVAE can be written as:

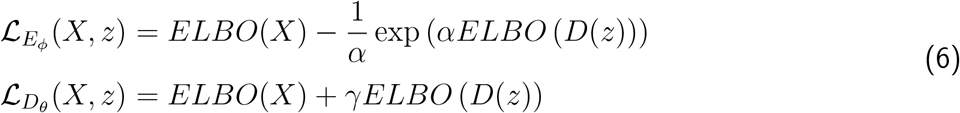

where *X* = {*x*_*i*_ | *i*^*th*^modality}, and *α >*= 0, *γ >*= 0 are hyperparameters. *ELBO*(*X*) is defined in Equation (1). *ELBO*(*D*(*z*)) can be defined similarly, but with *D*_*θ*_(*z*)_*i*_ = *p*_*θ*_ (*x*_*i*_ | *z*) replacing x in Equation (1). Following ELBO(*D*(*z*) in Equation 1, ELBO(*D*(*z*) can be written as:

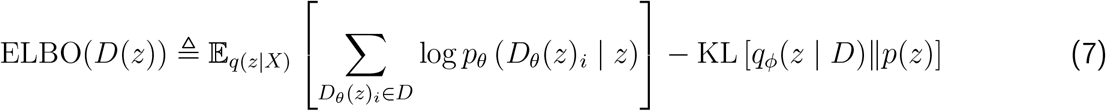

Equation (6) represents a min-max game between the encoders and the decoders. The encoders are encouraged, via the ELBO value, to differentiate between real samples (high ELBO) and generated samples (low ELBO), while the decoders aim to generate samples that “fool” the encoders (Figure 1).

**Figure 1:**
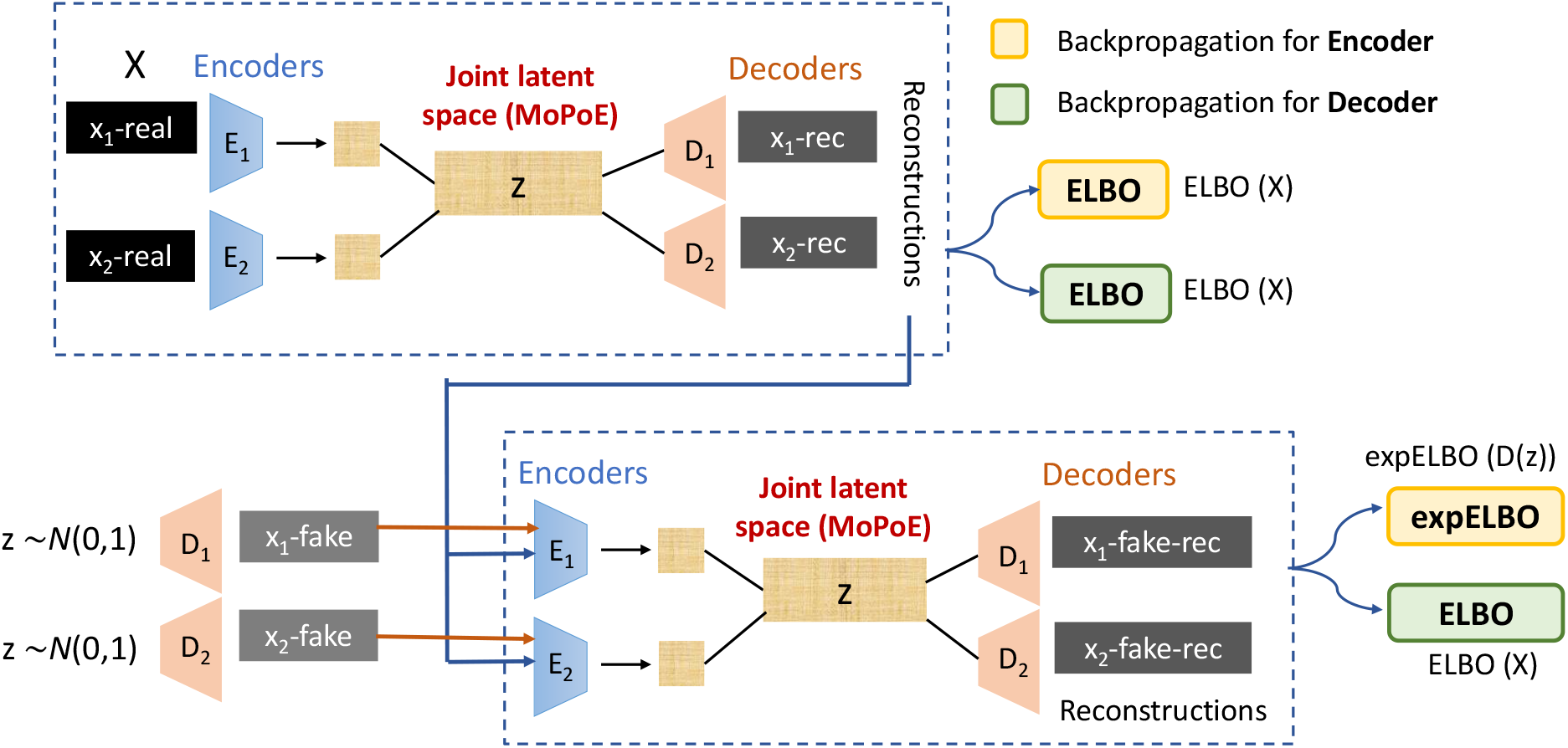
Training flow of mmSIVAE. The ELBO for real samples is optimized for both encoders and decoders, while the encoders also optimize the expELBO to ‘push away’ generated samples from the latent space. The decoders optimize the ELBO for the generated samples to ‘fool’ the encoders.

### 3.1 Theoretical analysis

In this section, we extend the theoretical analysis of SIVAE (Daniel and Tamar, 2021) for multimodal data and analyze the Nash equilibrium of the min-max game shown in Equation (6). Here, the encoder is represented by the approximate joint posterior distirbution *q* = *q*(*z*|*X*) and the decoder is represented by *d* = *p*_*d*_(*X*|*z*). *p*_*data*_(*X*) represents the data distribution and *p*(*z*) denote a prior distribution over the latent variable z. Using the conditional indepence of the modalities and the linearity of the expectation, we can denote *p*_*d*_(*x*_*i*_) = 𝔼_*p*(*z*)_ [*p*_*d*_(*x*_*i*_ | *z*)], which is the distribution of generated samples.

We can further denote *ELBO*(*X*) and *ELBO*(*D*(*z*)) from Equation (6) as *W* (*X*; *d, q*) and *W* (*D*; *d, q*) respectively, as shown below. *D* is the collection of decoders *D* = {*D*_*θ*_(*z*)_*i*_ | *i*^*th*^decoder}.

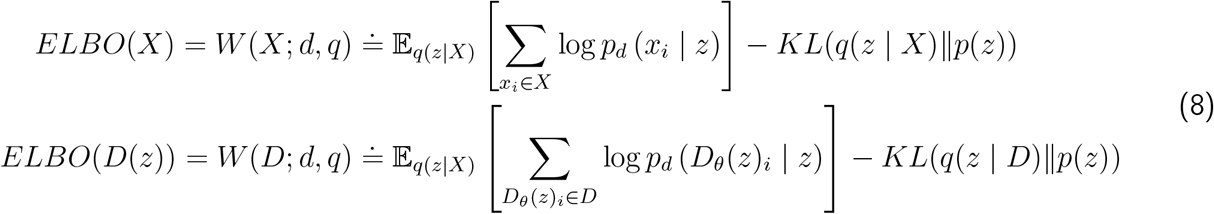

We consider a nonparametric setting where d and q can be any distribution. For some z, let D(z) be a multimodal sample from *p*_*d*_(*X* | *z*) The objective functions for q and d are given by:

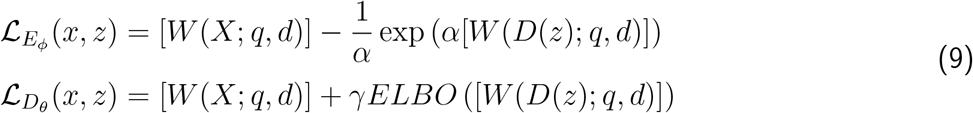

The complete mmSIVAE objective functions considers the expectation of the encoder and decoder losses over real (*p*_*data*_) and generated (*p*_*d*_) samples.

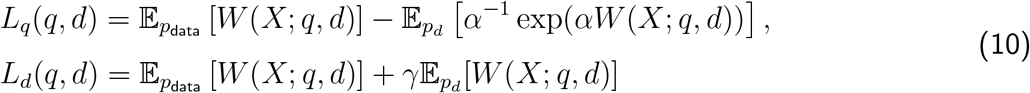

A Nash equilibrium point (*q*^*^, *d*^*^) satisfies *L*_*q*_ (*q*^*^, *d*^*^) ≥ *L*_*q*_ (*q, d*^*^) and *L*_*d*_ (*q*^*^, *d*^*^) ≥ *L*_*d*_ (*q*^*^, *d*) ∀*q, d*.

Given a *d*, let *q*^*^(*d*) satisfy *L*_*q*_(*q*^*^(*d*), *d*) ≥ *L*_*q*_(*q, d*) ∀ *q*.

#### Lemma 1

If 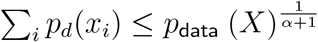 for all *X* for which (*X*) > 0,we have that *q*^*^(*d*) satisfies *q*^*^(*d*)(*z*|*X*) = *p*_*d*_(*z*|*X*), and *W* (*X*; *q*^*^(*d*), *d*) = ∑_*i*_ log *p*_*d*_(*x*_*i*_).

#### Lemma 2

Let *d*^*^ be defined so that 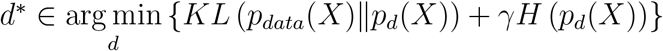 {*KL* (*p*_*data*_(*X*)∥*p*_*d*_(*X*)) + *γH* (*p*_*d*_(*X*))}, where *H*(·) is the Shannon entropy of the generated data samples. If *q*^*^ = *p*_*d*_^*^ (*z* | *x*), (*q*^*^, *d*^*^) is a Nash equilibrium of Equation (10).

The proofs of Lemma 1 and 2 are provided in Appendix A.

### 3.2 Modeling of the joint posterior - MOPOE

In Section 2.3, we discuss the limitations of commonly used techniques POE and MOE for aggregating information from multiple modalities in the latent space. In this work, we adopted a generalization of both MoE and PoE, called Mixture-of-Product-of-Experts (MoPoE) as shown below.

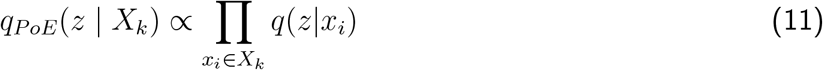

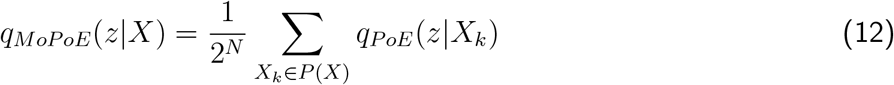

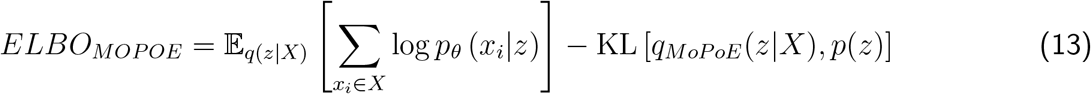

Here, *X*_*k*_ denotes a random subset of the N modalities, and *P* (*X*) signifies the power set of all N modalities. MoPoE can be viewed as a hierarchical distribution: first, the unimodal posterior approximations for a subset *X*_*k*_ are merged via PoE, followed by the aggregation of subset approximations *q*_*ϕ*_(*z* | *X*_*k*_) through MoE. This approach leverages the advantages of both MoE and PoE, addressing their respective limitations.

#### Lemma 3

MoPoE generates a generalized ELBO with PoE and MoE being special cases.

To prove this lemma, first we need to show that ELBO_*MoPoE*_(*X*) is a valid multimodal ELBO, i.e. log *p*_*θ*_(*x*) ≥ ELBO_*MoPoE*_(*X*). Next we will show that MOPOE is a generalized case of both POE and MOE. The proofs have been provided in Appendix B.

### 3.3 Model training and hyperparameters

Algorithm 1 shows the detailed training procedure of mmSIVAE. Expanding mmSIVAE’s objective function, which is minimized from Equation 6 with the complete set of hyperparameters.

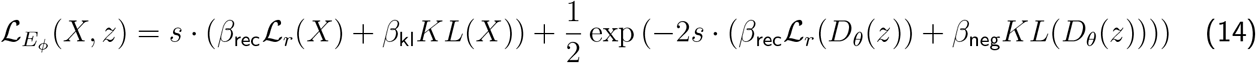

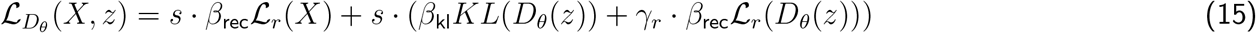

where 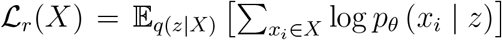 denotes the summation of reconstruction error of real samples X across the 2 modalities. *KL*(*X*) = KL [*q*(*z* | *X*)∥*p*(*z*)] denotes the KL divergence between the joint posterior and prior *p*(*z*).

Similarly, 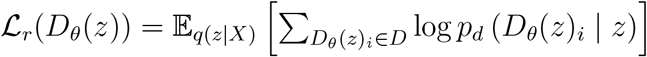 denote the summation of the reconstruction error of the generated samples *D*(*z*) across the 2 modalities. *KL*(*D*(*z*)) = *KL*(*q*(*z* | *D*)∥*p*(*z*) denote the KL divergence between the joint posterior of generated samples *D*(*z*) and the prior *p*(*z*).

The hyperparameters *β*_rec_ and *β*_kl_ control the balance between inference and sampling quality respectively. When *β*_rec_ *> β*_kl_, the optimization is focused on good reconstructions, which may lead to less variability in the generated samples, as latent posteriors are allowed to be very different from the prior. When *β*_rec_ *< β*_kl_, there will be more varied samples, but reconstruction quality will degrade.

Each ELBO term in Equations 14 and 15 can be considered as an instance of *β*_VAE_ and can have different *β*_rec_ and *β*_kl_ parameters. However, we set them all to be the same, except for the ELBO inside the exponent in Equation 14. For this term, *β*_kl_ controls the repulsion force of the posterior for generated samples from the prior. We denote this specific parameter as *β*_neg_.

As reported in the SIVAE paper Daniel and Tamar, 2021, the decoder tries to minimize the reconstruction error for generated data, which may slow down convergence, as at the beginning of the optimization the generated samples are of low quality. *γ*_*r*_ is a hyperparameter that multiplies only the reconstruction term of the generated data in the ELBO term of the decoder in Equation 15.

#### Algorithm 1 Training multimodal soft-introVAE (mmSIVAE)

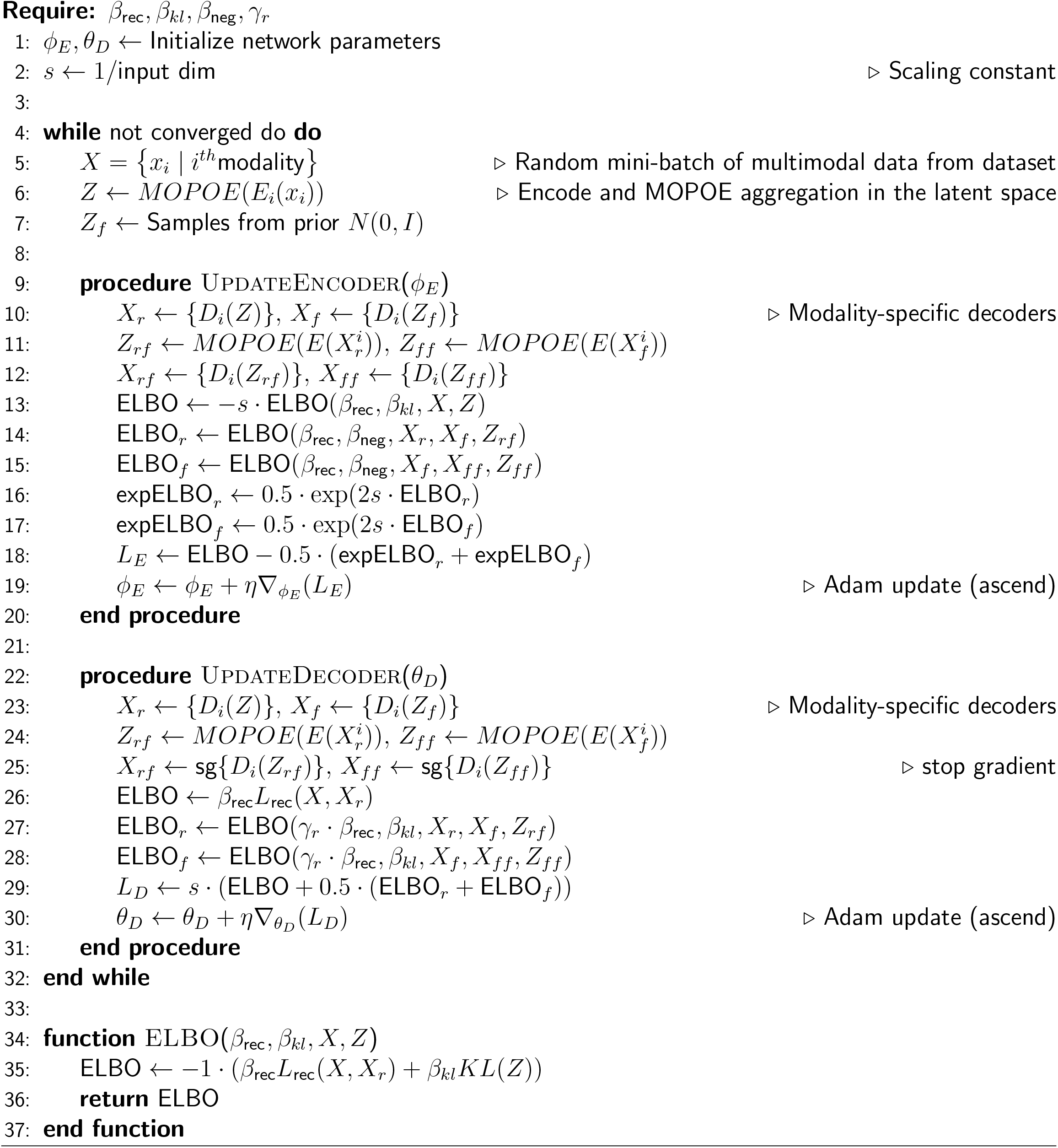

Finally, we use a scaling constant s to balance between the ELBO and the expELBO terms in the loss, and we set s to be the inverse of the input dimensions Daniel and Tamar, 2021. This scaling constant prevents the expELBO from vanishing for high-dimensional input in Equation 14.

### 3.4 Multimodal normative modeling

mmSIVAE (Figure 1) has separate encoders to encode the 2 modalities into their corresponding latent parameters (mean and variance). The unimodal latents were aggregated through the MoPoE approach to estimate the shared latent parameters (joint latent distribution). The shared latents were passed through the modality-specific decoders to reconstruct each modality. The model was first trained to characterize the healthy population cohort. We assumed disease abnormality can be quantified by measuring how AD subjects deviate from the joint space (latent deviations) Lawry Aguila et al., 2022, 2023 or from the reconstruction errors of healthy controls (feature-space deviations) Kumar, Earnest, et al., 2024; Kumar, Payne, and Sotiras, 2023. During inference, the trained model was applied on the AD cohort to estimate both latent and feature deviations.

#### Patient-specific deviations - latent and feature space

##### Mahalanobis distance

To quantify how much AD each subject deviates from the latent distribution of healthy controls, we measured the Mahalanobis distance Lawry et al., 2023, which accounts for correlations between latent vectors.

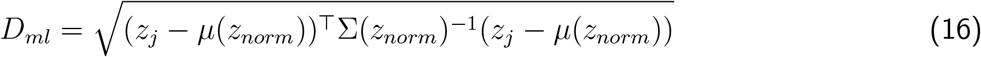

where *z*_*j*_ ≡ *q*(*z*_*j*_|*X*_*j*_) is a sample from the joint posterior distribution for subject *j. µ*(*z*_*norm*_) and ∑(*z*_*norm*_) represent the mean and covariance of the healthy cohort latent position. We additionally derived a multivariate feature-space deviation index based on the Mahalanobis distance that quantifies how the reconstruction error of an AD subject deviate from the reconstruction errors of controls.

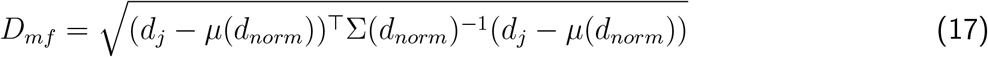

where 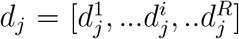 is the mean squared reconstruction error between original and reconstructed input for subject j and brain region *i* = [1, 2, ..*R*]. *µ*(*d*_*norm*_) and *σ*(*d*_*norm*_) are the mean and covariance of the healthy cohort reconstruction error respectively.

##### Z-score latent and feature deviations

To analyze which latent dimensions and brain regions are associated with abnormal deviations due to AD pathology, we calculated both latent space Z-scores *Z*_*ml*_ (for each latent dimension) and feature-space Z-scores *Z*_*mf*_ . *z*_*ij*_ and *d*_*ij*_ represent the latent values and reconstruction error of test subject j for i-th position respectively. 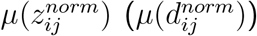 and 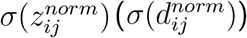 are the mean and standard deviations of healthy cohort latent values (reconstruction error) respectively.

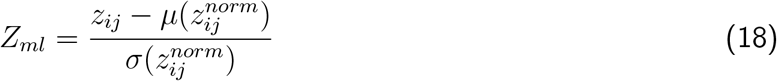

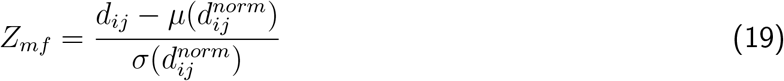

## 4 Experimental set-up

### 4.1 Data and feature preprocessing

For training, we selected 248 cognitively unimpaired (healthy control) subjects from the Alzheimer’s Disease Neuroimaging (ADNI) dataset Mueller et al., 2005 with Clinical Dementia Rating (CDR) = 0 and no amyloid pathology. We used regional brain volumes extracted from T1-weighted MRI scans and regional Standardized Uptake Value Ratio (SUVR) values extracted from AV45 Amyloid PET scans as the two input modalities for our model (Figure 1). Both brain volumes and SUVR values were extracted from 66 cortical (Desikan-Killiany atlas) and 24 subcortical regions. For model evaluation, we used 48 healthy controls for a separate holdout cohort and a disease cohort of 726 amyloid positive AD spectrum (ADS) individuals across the following AD stages: preclinical stage with no symptoms (*CDR* = 0, *A*+)(*N* = 305), (b) *CDR* = 0.5(*N* = 236) and (c) *CDR >*= 1(*N* = 185).

Each brain ROI was normalised by removing the mean and dividing by the standard deviation of the healthy control cohort brain regions. We conditioned our model on the age and sex of patients, represented as one-hot encoding vectors, to remove the effects of covariates from the MRI and PET features. Details about ADNI data acquisition and preprocessing for T1-weighted MRI and amyloid AV45-PET are provided in Appendix B.

### 4.2 Baselines and implementation details

#### Baselines

Our proposed mmSIVAE was compared with the following baselines as follows: (i) Unimodal Soft-Instrospective VAE (SIVAE) Daniel and Tamar, 2021(Multimodal VAE (Aim 2B; mmVAE) Kumar, Payne, and Sotiras, 2023, (ii) unimodal VAE Pinaya et al., 2021 and **state-of-the-art multimodal VAE models**: mmJSD Sutter et al., 2020, JMVAE Suzuki et al., 2016 and MVTCAE Hwang et al., 2021. For multimodal methods (mmSIVAE and mmVAE), model performance using different aggregation techniques (POE, MOE and MOPOE) were compared. Unimodal methods (SIVAE and VAE) had a single encoder and decoder and used either a single modality (MRI/amyloid) or both modalities as a single concatenated input. Models mmJSD, JMVAE and MVTCAE were implemented using the open-source Multi-view-AE python package Aguila et al., 2023.

#### Model hyperparameters

The range of the search was [0.05, 1.0] for *β*_kl_ and *β*_rec_ and [*β*_kl_, 5*β*_kl_] for *β*_neg_. Following a hyperparameter tuning by grid search, the final values selected are *β*_kl_ = *β*_kl_ = 1 and *β*_neg_ = 10. As followed in the SIVAE paper, we set constant *γ*_*r*_ = 10^−8^ and s = 1/number of features = 1/90.

All models were implemented in Pytorch and trained using Adam optimizer with hyperparameters as follows: epochs = 500, learning rate = 10^−5^, batch size = 64 and latent dimensions in the range [5,10,15,20]. The encoder and decoder networks have 2 fully-connected layers of sizes 64, 32 and 32, 64 respectively.

## 5 Results and Discussion

### 5.1 Reconstruction of healthy controls

Our first objective was to evaluate whether the introspective VAE (mmSIVAE) achieved better reconstruction performance for cognitively unimpaired healthy controls compared to a standard vanilla VAE. Furthermore, we examined whether multimodal methods, which integrate information from multiple modalities in the latent space, demonstrated improved reconstruction accuracy compared to their unimodal counterparts. To address this, we visualized the average reconstruction errors for each cortical and subcortical region across our proposed mmSIVAE and baseline models, including SIVAE, multimodal VAE (mmVAE; Aim 2B), and vanilla VAE (Figure 3). For a fair comparison, both mmSIVAE and mmVAE employed the MOPOE technique to aggregate multimodal information within the latent space. In contrast, the unimodal methods, SIVAE and vanilla VAE, utilized a single encoder and decoder, with both modalities (MRI and amyloid) concatenated as a single unimodal input. The brain atlases showing the reconstruction errors for both MRI volumes and amyloid SUVR were visualized using the ggseg package in Python.

Across all brain regions, mmSIVAE demonstrated the lowest reconstruction error for both modalities when compared to the baseline models (Figure 3). Introspective VAE methods, such as mmSIVAE and SIVAE, outperformed vanilla VAE approaches (mmVAE and VAE) in reconstruction accuracy. Additionally, multimodal methods (mmSIVAE and mmVAE), which integrate information from multiple modalities, exhibited lower reconstruction errors than unimodal VAEs that treated multiple modalities as a single unimodal input (Figure 3).

### 5.2 Evaluating outlier detection performance

The latent mahalanobis deviations *D*_*ml*_ (Equation 16) and the feature mahalanobis deviations *D*_*mf*_ (Equation 17) were estimated independently for the ADS cohort and separate holdout cohort of healthy controls with no cognitive impairment. For both deviation metrics, subjects with statistically significant (*p <* 0.001) deviations Tabachnick et al., 2013 were labeled as abnormal/outliers. A good normative model is supposed to correctly identify disease subjects as outliers and healthy individuals within the normative distribution. Following previous work, we used the positive likelihood ratio to assess how well our mmSIVAE model can detect outliers.

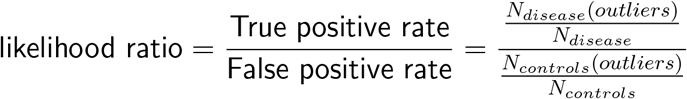

Latent deviations (*D*_*ml*_) consistently achieved higher likelihood ratios compared to feature space deviations (*D*_*mf*_), as shown in Tables 1 and 2. This demonstrates that calculating deviations within the multimodal joint latent space enhances normative performance and improves outlier detection compared to regional space deviations. Among the models, mmSIVAE with MOPOE latent space aggregation achieved the highest likelihood ratios, outperforming all baselines. Introspective VAE methods (mmSIVAE and SIVAE) exhibited superior normative performance compared to vanilla VAE-based methods (mmVAE and unimodal VAE). Furthermore, in both mmSIVAE and mmVAE, MOPOE aggregation proved to be more effective for outlier detection than POE or MOE, validating its selection as the preferred aggregation strategy by taking advantage of the strengths of both approaches. In general, increasing the number of latent dimensions to 15 or 20 resulted in higher likelihood ratios, further improving performance.

**Table 1:**
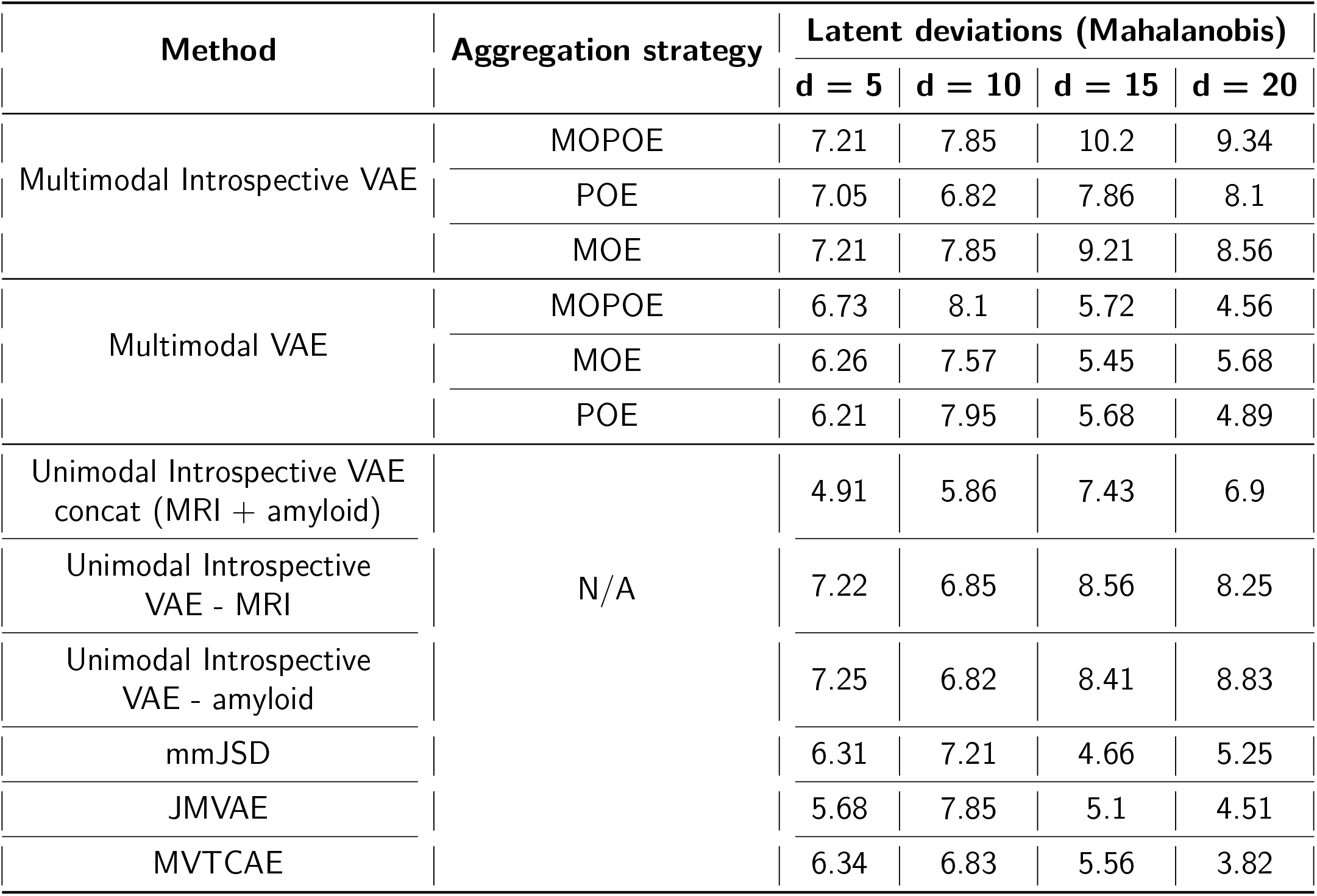
Likelihood ratio corresponding to outliers derived from *D*_*ml*_ for different latent dimensions d = 5, 10, 15 and 20. Higher likelihood ratio indicate better detection of outlier AD patients.

**Table 2:** Likelihood ratio corresponding to outliers derived from *D*_*mf*_ for different latent dimensions d = 5, 10, 15 and 20. Higher likelihood ratio indicate better detection of outlier AD patients.

| Method | Aggregation strategy | Feature deviations (Mahalanobis) |  |  |  |
| --- | --- | --- | --- | --- | --- |
| | | $d = 5$ | $d = 10$ | $d = 15$ | $d = 20$ |
| Multimodal Introspective VAE | MOPOE | 2.42 | 2.76 | 3.29 | 3.34 |
|  | POE | 2.57 | 2.68 | 2.85 | 3.09 |
|  | MOE | 2.45 | 2.77 | 2.55 | 2.94 |
| Multimodal VAE | MOPOE | 1.85 | 2.52 | 2.16 | 1.76 |
|  | MOE | 1.67 | 1.92 | 2.22 | 1.32 |
|  | POE | 1.37 | 1.53 | 1.77 | 1.68 |
| Unimodal Introspective VAE<br>concat (MRI + amyloid) | N/A | 1.96 | 2.04 | 2.62 | 2.85 |
| Unimodal Introspective<br>VAE - MRI |  | 2.21 | 2.45 | 2.72 | 3.05 |
| Unimodal Introspective<br>VAE - amyloid |  | 2.32 | 2.57 | 2.75 | 2.91 |
| mmJSD |  | 2.21 | 1.81 | 1.4 | 1.25 |
| JMVAE |  | 1.68 | 2.33 | 2.1 | 1.75 |
| MVTCAE |  | 1.35 | 1.68 | 1.52 | 1.21 |

A similar trend is observed for likelihood ratios generated by feature-space Mahalanobis deviations (*D*_*mf*_), as shown in Table 2. Our proposed mmSIVAE achieved the greatest separation between control and disease cohorts, evidenced by the highest Earth Mover’s Distance (34.043) between the *D*_*mf*_ distributions of the two cohorts (Figure 4). Introspective VAE-based methods (mmSIVAE and SIVAE) demonstrated more pronounced separation (reduced overlap and greater distance) between control and disease cohorts compared to VAE-based methods (mmVAE and VAE), as illustrated in Figure 4.

### 5.3 Association with cognitive performance

We demonstrated the clinical validation of the multimodal latent deviations *D*_*ml*_ generated by our proposed mmSIVAE with MoPoE aggregation by assessing the association of *D*_*ml*_ wih cognitive decline. Towards this end, we examined the association between *D*_*ml*_ and 4 neuropsychological composite cognitive functions : memory, executive functioning, language and visual-spatial. These neuropsychological composites represent different domains of cognitive and functional performance. Details of how these scores were extracted from the ADNI dataset are listed in Appendix B.2.4. The correlations between *D*_*ml*_ and the composites were estimated using the Pearson correlation coefficient with Steiger’s Z-test Steiger, 1980 being used to compare the correlations between our proposed mmSIVAE and other baselines (unimodal SIVAE, mmVAE and unimodal VAE). For fair comparison, all multimodal methods (mmSIVAE and mmVAE) aggregated multiple modalities using MOPOE. Steiger’s Z-test is used to compare dependent correlations, which means correlations that share at least one variable and are calculated from the same dataset. It helps determine if there is a significant difference between two correlations.

Our proposed multimodal soft-introspective VAE (mmSIVAE) showed higher correlation (in magnitude) with impairment (lower scores) in memory, executive, language and visual-spatial skills compared to the baselines (Table 3). The correlation between cognitive decline and latent deviations of mmSIVAE differs significantly from the correlations between cognitive decline and latent deviations of mmVAE and VAE. However, there are no significant differences in the correlations with cognitive compounds between mmSIVAE and SIVAE (Table 3).

**Table 3:** Pearson correlation coefficients to show the correlation between *D*_*ml*_ of different methods with the neuropsychological composites : memory, executive, language and visual-spatial. The numbers in () represent the Steiger’s Z-test Steiger, 1980 p values to compare the correlations between mmSIVAE and the baselines. p-values more than 0.05 (marked in red) indicate that the differences are not significant. For fair comparison, all multimodal methods used MOPOE aggregation.

| Method | Memory | Executive | Language | Visual-spatial |
| --- | --- | --- | --- | --- |
|  | Pearson correlation coefficient (Sigel’s Z-test p-value) |  |  |  |
| Multimodal introVAE | -0.573 | -0.467 | -0.421 | -0.442 |
| Unimodal introVAE – concat (MRI + amyloid) | -0.521 (0.304) | -0.472 (0.903) | -0.375 (0.064) | -0.326 (0.032) |
| Multimodal VAE | -0.423 (0.0087) | -0.385 (0.027) | -0.392 (0.174) | -0.294 (0.011) |
| Unimodal VAE – concat (MRI + amyloid) | -0.394 (0.0023) | -0.402 (0.074) | -0.374 (0.071) | -0.306 (0.018) |

### 5.4 Interpretability analysis

#### 5.4.1 Mapping from latent to feature deviations

Ideally, all latent dimensions can be used to reconstruct the input data and quantify feature-space deviations. The latent dimensions with statistically significant (*p <* 0.05) mean absolute Z-scores *Z*_*ml*_ (Equation 18) indicate the latent dimensions which show deviation between control and disease cohorts. This can provide an interpretation of how latent space deviations can be mapped to deviations in the feature-space. We passed these selected latent vectors through the decoders setting the remaining latent dimensions and covariates to be 0 such that the reconstructions and feature deviations *Z*_*mf*_ (Equation 19) reflect only the information encoded in the selected latent vectors. From Table 1, we observed mmSIVAE with MOPOE latent space aggregation had the maximum likelihood ratio for d = 15 (number of latent dimensions). We identified 5 out of 15 latent dimensions (6,7,8,14 and 15) whose mean absolute *Z*_*ml*_ *>* 1.96 (*p <* 0.05) and used them for generating the feature-space deviations *Z*_*mf*_ (Figure 5).

#### 5.4.2 Brain regions associated with AD abnormality

We visualized the pairwise differences in the magnitude of abnormal deviations (*Z*_*mf*_) in each region between amyloid negative CU individuals and disease groups along the ADS: (i) CDR = 0 (preclinical AD), (ii) CDR = 0.5 (very mild dementia), and (iii) CDR ¿= 1 (mild or more severe dementia). Our aim was to validate the derived regional abnormal deviations by examine examining whether these deviations showed increased group differences across progressive CDR stages. We quantified group differences using Cohen’s d-statistic effect size, calculated separately for each modality. A higher effect size when comparing regional MRI volumes indicated lower gray matter volume (more atrophy). Similarly, a higher effect size when comparing amyloid or tau uptake indicated elevated SUVR uptake (higher amyloid and tau loads) compared to the amyloid-negative CU group.

Region-level group differences in MRI atrophy were most evident within the temporal, parietal and hippocampal regions which is consistent with the observations in existing literature Leech et al., 2014; Young et al., 2014 (Figure 2B). Higher group differences in amyloid loading were mostly observed in the accumbens, precuneus, frontal and temporal regions, which are sensitive to amyloid pathology accumulation Levitis et al., 2022 (Figure 6).

**Figure 2:**
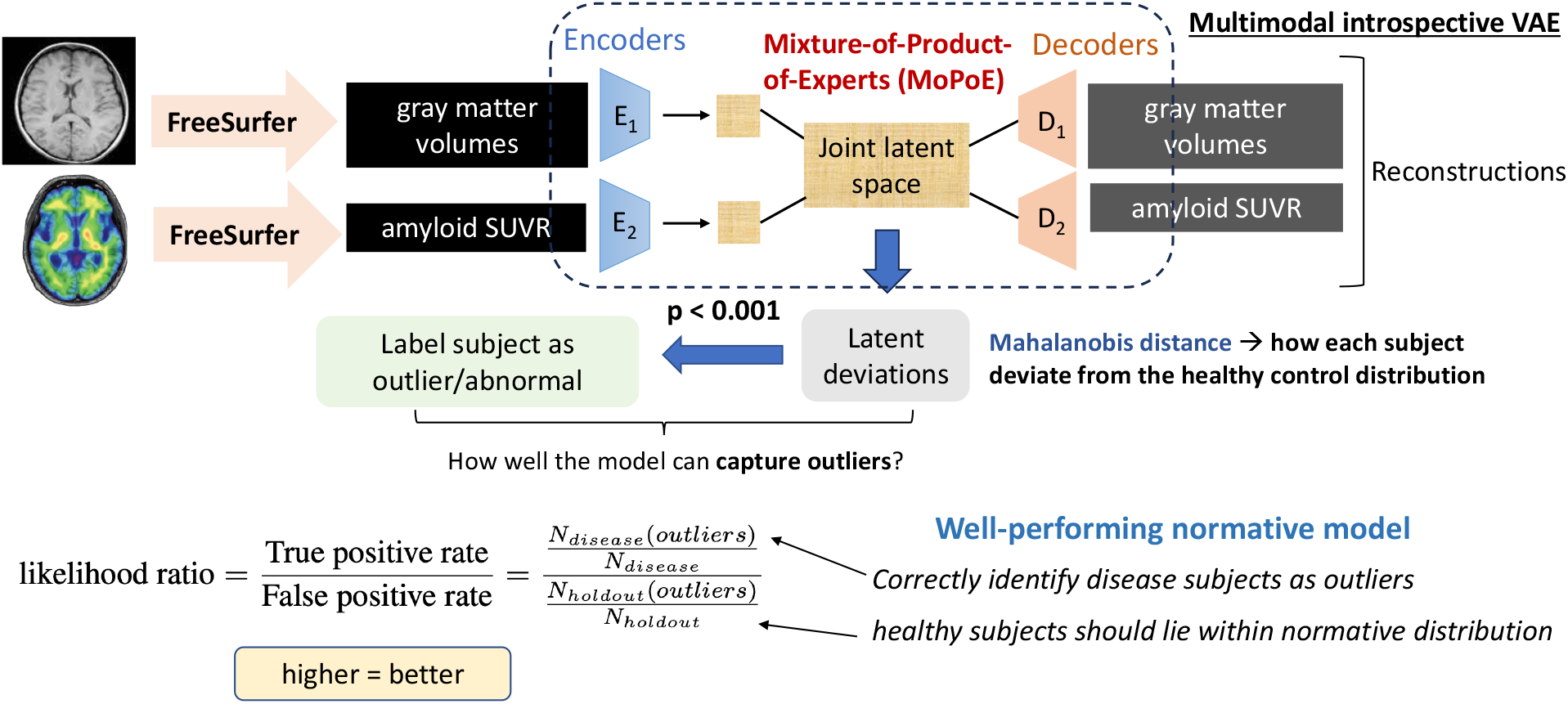
Normative modeling using multimodal soft-intro VAE (mmSIVAE). Deviations calculated in the latent space using Mahalonobis distance can be used to detect outlier AD patients. The normative performance is evaluated by the likelihood ratio (higher = better).

**Figure 3:**
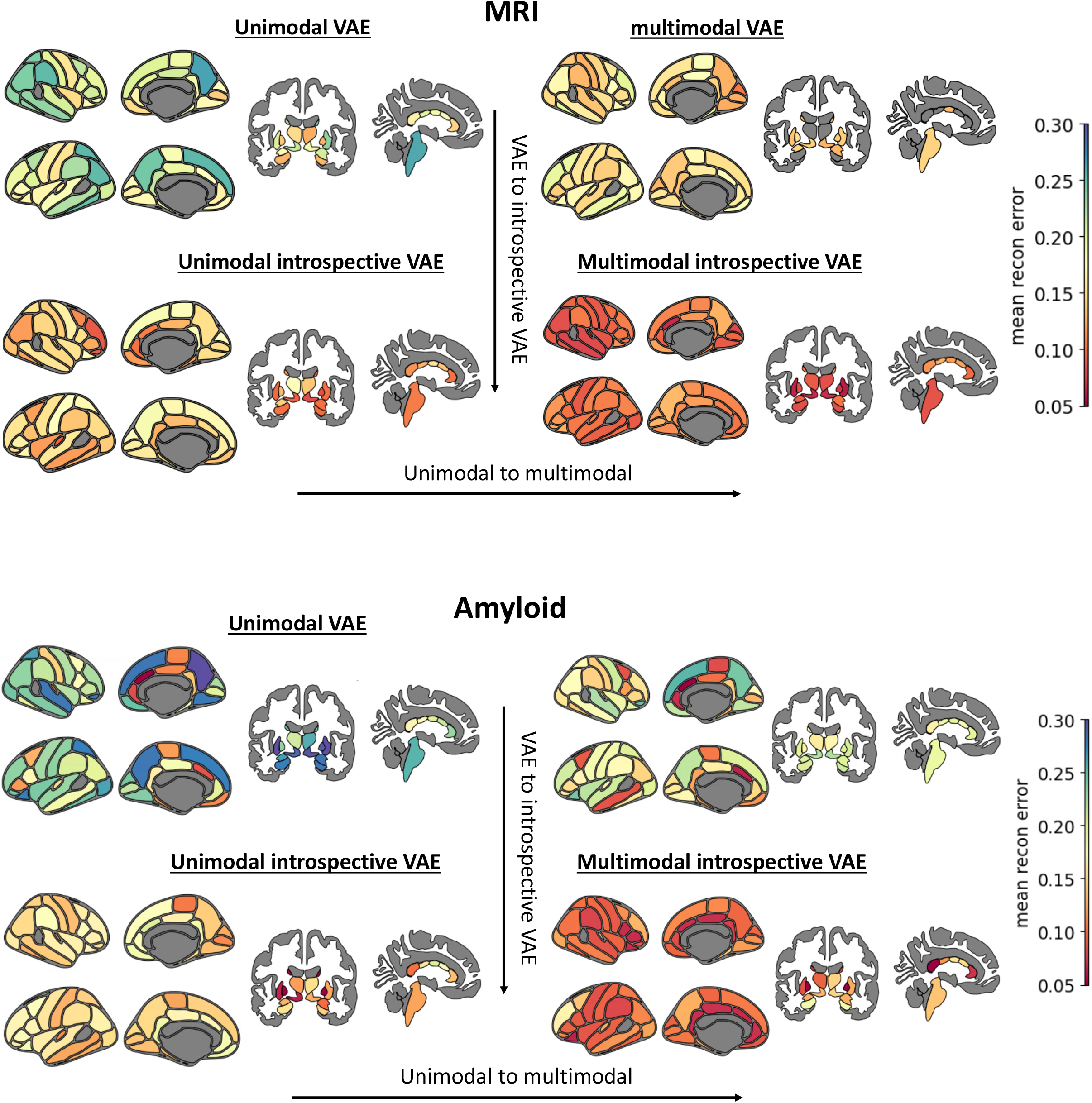
Average reconstruction error when reconstructing MRI volumes (top panel) and amyloid SUVR (bottom panel) for our proposed mmSIVAE and baselines (SIVAE, mmVAE and unimodal VAE).

**Figure 4:**
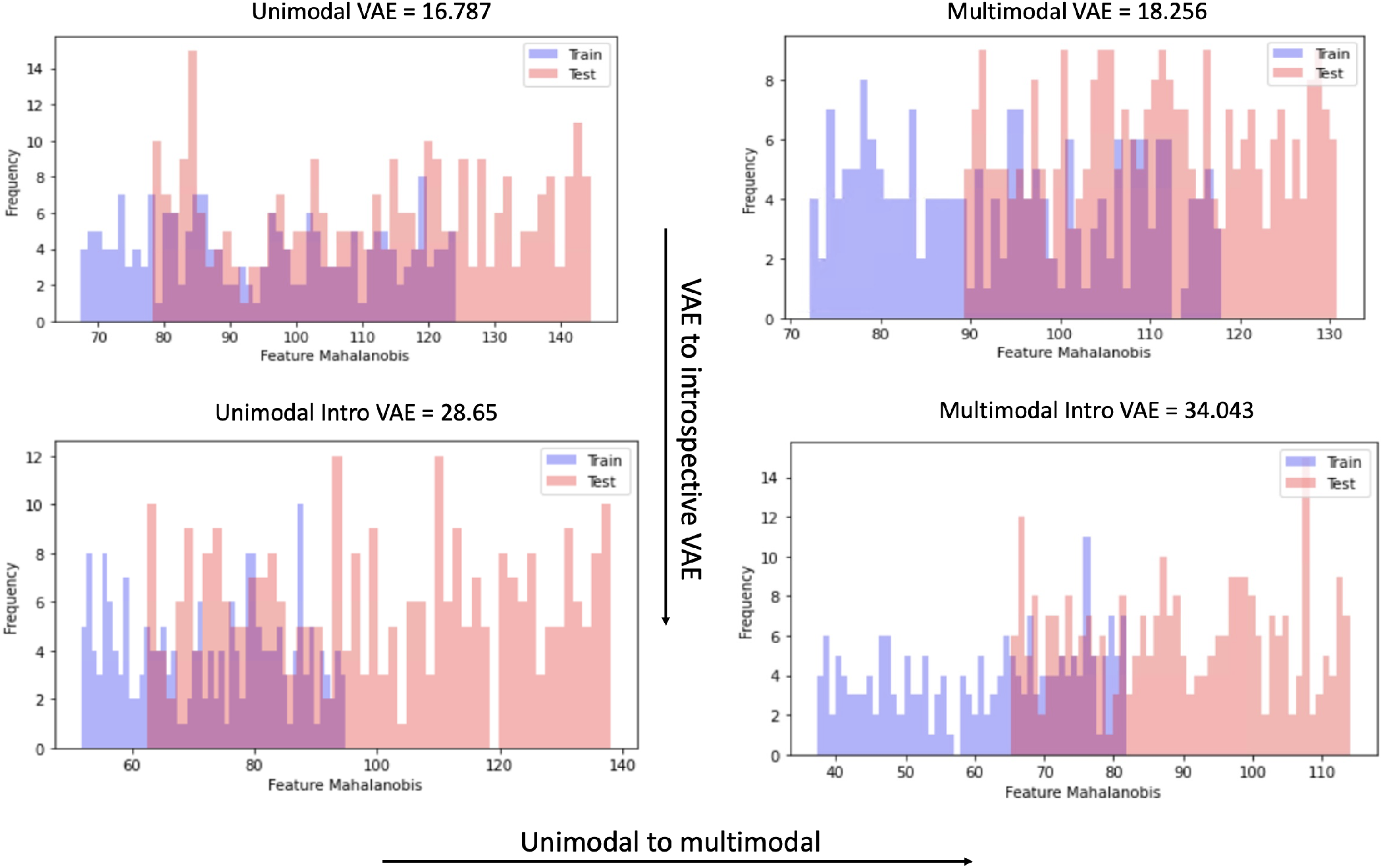
Histograms showing the distribution of feature mahalanobis deviations *D*_*mf*_ (Equation 17) for the test (AD patients) and a holdout healthy control cohort. The values in the caption of each subfigure indicate the Earth mover’s distance between the 2 distributions train and test. Higher distance indicate lesser overlap and better separation between the control and disease cohorts.

**Figure 5:**
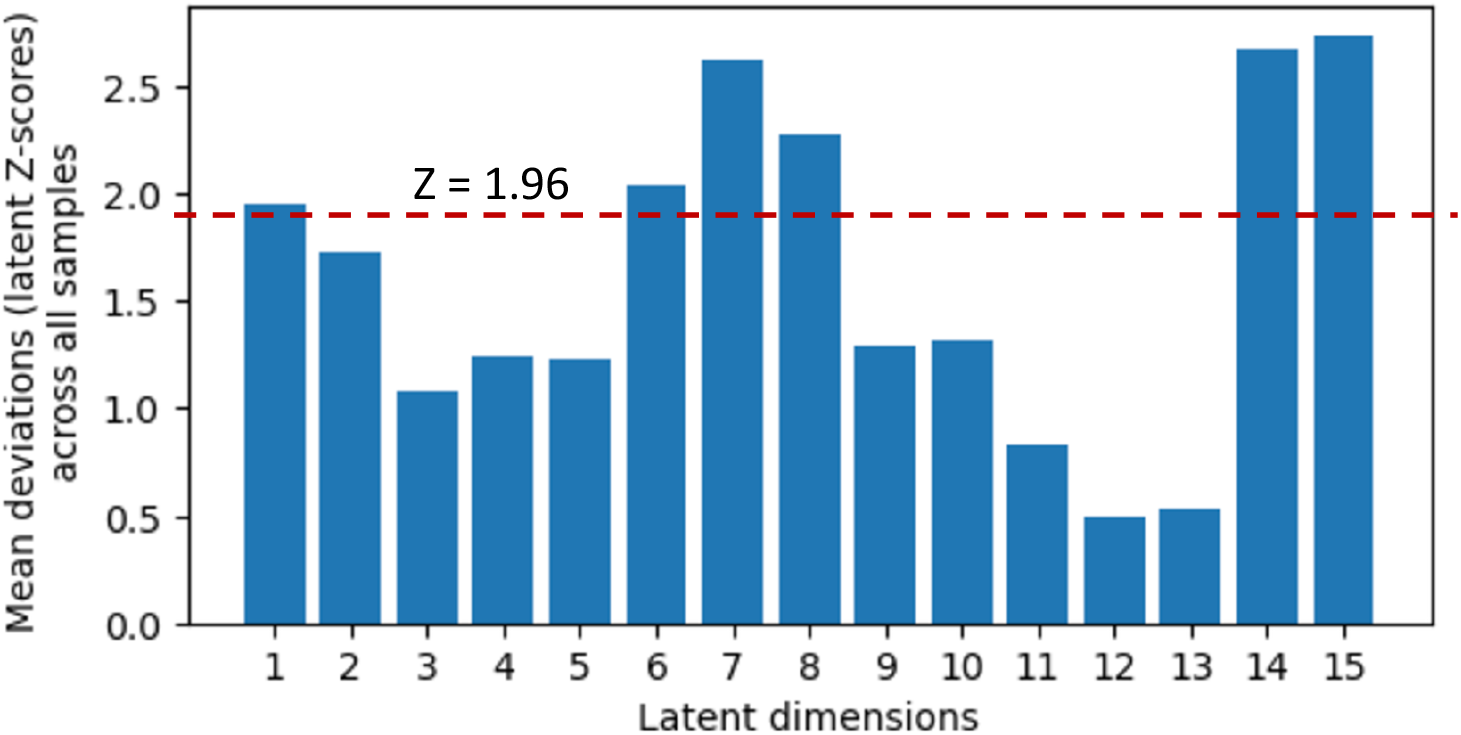
5 latent dimensions (6,7,8, 14 and 15) out of 15 with statistically significant deviations (mean absolute *Z*_*ml*_ *>* 1.96 or *p <* 0.05). The dotted red line indicates *Z >* 1.96. Latent dimensions above the dotted line were used for mapping to feature-space deviations.

**Figure 6:**
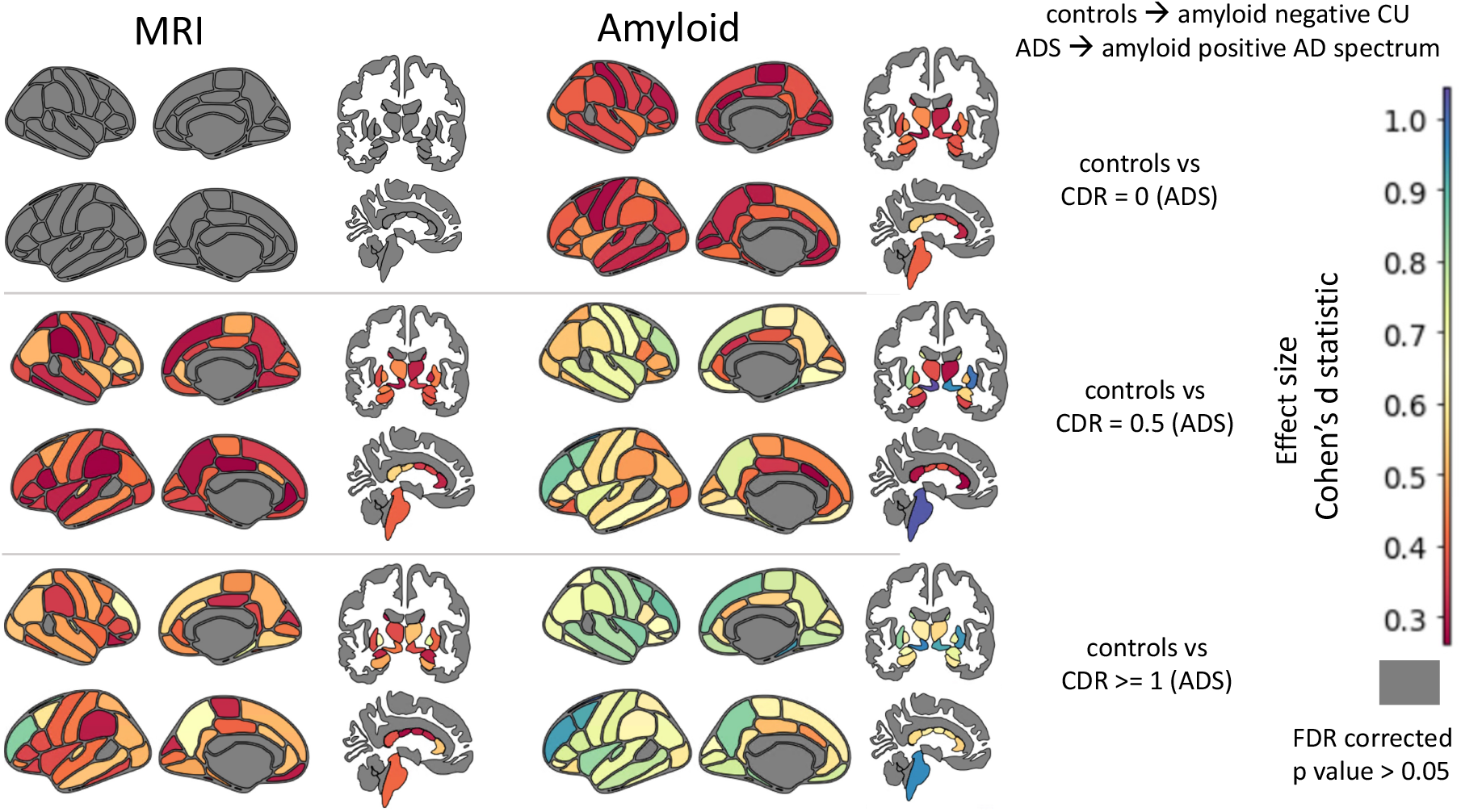
Brain atlas maps (Desikan-Killiany atlas for 66 cortical regions and Aseg atlas for 24 subcortical regions) showing the pairwise group differences in magnitude of deviations at each region between the amyloid negative CU group and each of the CDR groups. The color bar represents the effect size (Cohen’s d statistic). Effect sizes of d = 0.2, d = 0.5, and d = 0.8 are typically categorized as small, medium, and large, respectively. Gray regions represent the regions with no statistically significant deviations after FDR correction.

## 6 Conclusion

This study addressed key limitations of VAE-based deep normative models by proposing the multimodal soft-introspective VAE (mmSIVAE), which integrates the MOPOE strategy to improve multimodal information aggregation and outlier detection. Through analysis on the ADNI dataset, mmSIVAE demonstrated superior reconstruction accuracy for healthy controls and outperformed baseline methods in detecting outlier patients. The model effectively leveraged complementary multimodal data, achieving higher likelihood ratios and enhanced Out-of-Distribution (OOD) detection. Furthermore, the identification of significant Z-score deviations in latent dimensions allowed for the mapping of these deviations to feature-space abnormalities, providing insights into brain regions associated with Alzheimer’s disease. This work highlights the potential of mmSIVAE for advancing multimodal normative modeling in clinical applications.

## A Nash equilibrium - mmSIVAE

We can denote ELBO *W* (*X*; *d, q*) and *W* (*D*; *d, q*) as

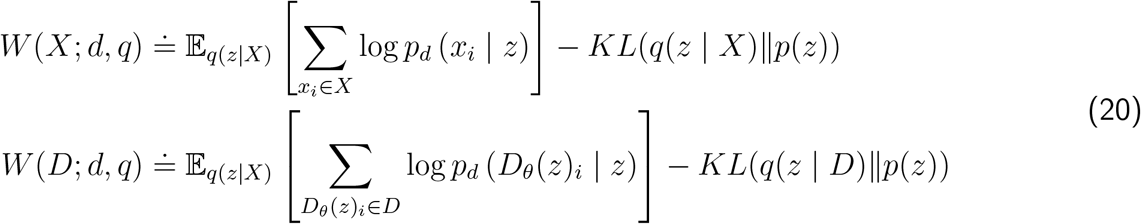

Here, the encoder is represented by the approximate joint posterior distribution *q* = *q*(*z*|*X*) and the decoder is represented by *d* = *p*_*d*_(*X*|*z*). The priors is *p*(*z*). Using the conditional independence of the modalities and the linearity of the expectation, we can denote *p*_*d*_(*x*_*i*_) = 𝔼_*p*(*z*)_ [*p*_*d*_(*x*_*i*_ | *z*)]. From the Radon-Nikodym Theorem, we have the following equalities:

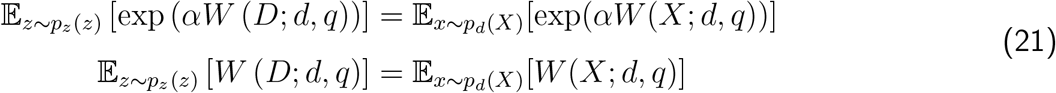

and the ELBO satisfies

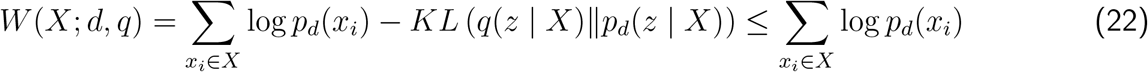

where 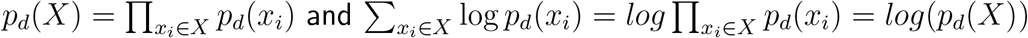

We consider a non-parametric setting where d and q can be any distribution. For some z, let D(z) be a multimodal sample from *p*_*d*_(*X* | *z*) The objective functions for q and d are given by:

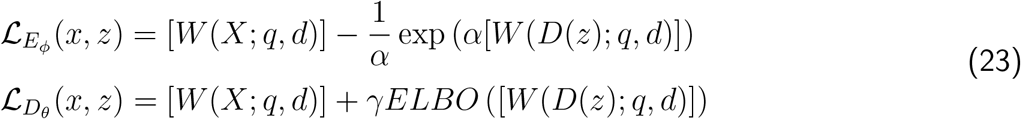

The complete multi-modal Soft-IntroVAE objective functions considers the expectation of the encoder and decoder losses over real and generate samples

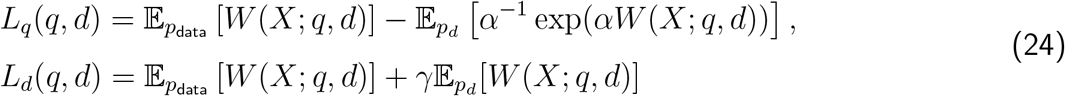

A Nash equilibrium point (*q*^*^, *d*^*^) satisfies

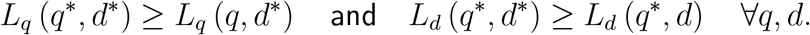

Given a *d*, let *q*^*^(*d*) satisfy *L*_*q*_(*q*^*^(*d*), *d*) ≥ *L*_*q*_(*q, d*) ∀ *q*.

### Lemma 1

If 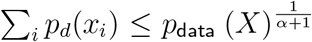 for all *X* for which *p*_data_ (*X*) *>* 0, we have that *q*^*^(*d*) satisfies *q*^*^(*d*)(*z*|*X*) = *p*_*d*_(*z*|*X*), and *W* (*X*; *q*^*^(*d*), *d*) = ∑_*i*_ log *p*_*d*_(*x*_*i*_).

### Proof of Lemma 1

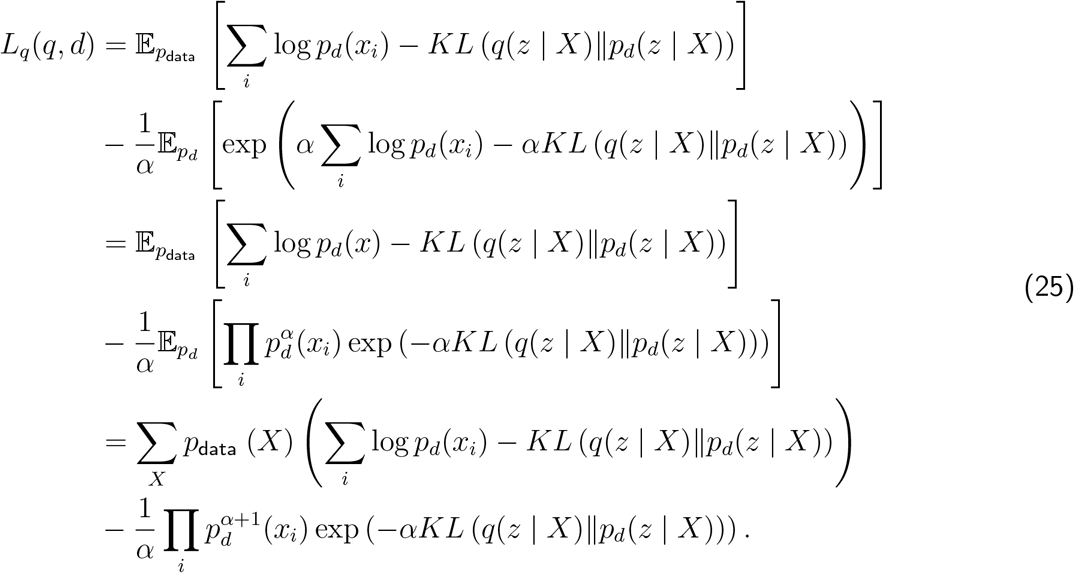

Consider a X for which *p*_*data*_(*X*) *>* 0. We have that *q*^*^(*d*(*z*|*X*) which maximizes

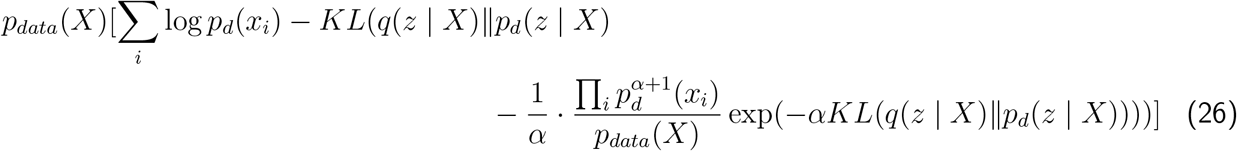

We consider the function 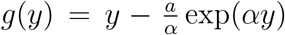 with *g*^*′*^(*y*) = 1 − *a* exp(*αy*) and a maximum 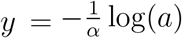. In the multi-modal Intro-SoftVAE formulation, 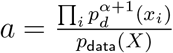 and *y* = −*KL* (*q*(*z* | *X*)∥*p* (*z* | *X*)) ≤ 0.

When *a >* 1, the maximum will be attained at 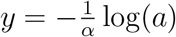. And if *a* ≤ 1, the maximum is attained at *y* = 0. For an *X* such that *p*_*data*_(*X*) = 0, *q*(*z*|*X*) is the maximizer of 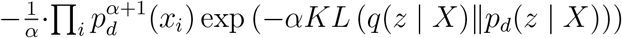

Since the KL divergence is always positive and 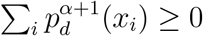, the maximum is obtained for −*KL* (*q*(*z* | *X*)∥*p*_*d*_(*z* |*X)) =* 0. Thus for every *X* the maximum is obtained for *KL* (*q*(*z* | *X*)∥*p*_*d*_(*z* | *X*)) = 0.

So we can see that *q*^*^(*d*)(*z*|*X*) = *p*_*d*_(*z*|*X*), and 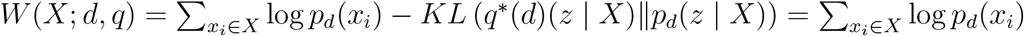

### Lemma 2

Let *d*^*^ be defined such that

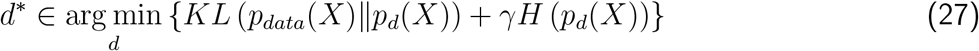

where *H*(·) is the Shannon entropy of the generated data samples. Let *q*^*^ = *p*_*d*_^*^ (*z* | *x*). Then (*q*^*^, *d*^*^) is the Nash equilibrium of min-max game between the encoders and decoders.

### Proof of Lemma 2

From Lemma 1, we have 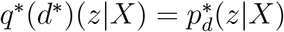. Let d be some decoder parameters (*p*_*d*_(*x* | *z*)).

From Equation 20, we have 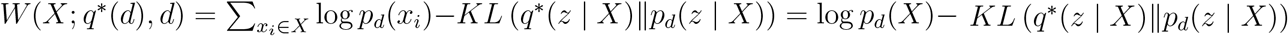.

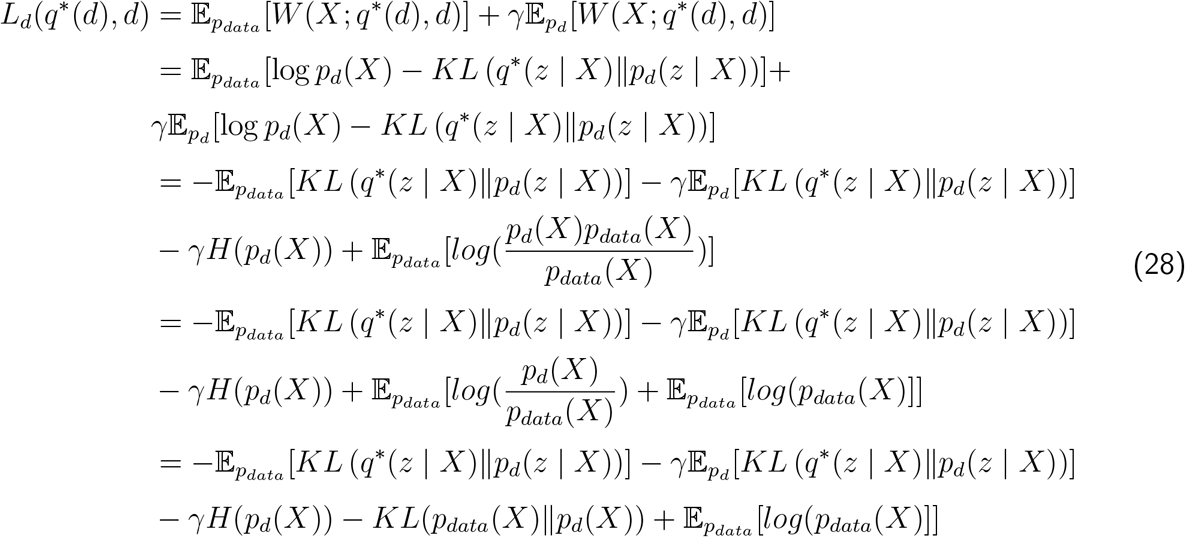

Since 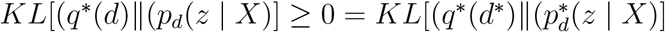 and 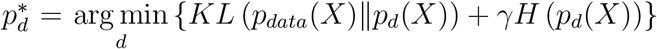, we have 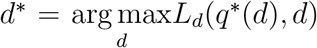. Also, since 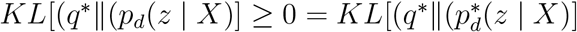, we have that 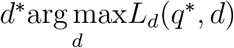.

Hence, from Lemma 1 and Lemma 2, we conclude that (*q*^*^, *d*^*^) is a Nash equilibrium of multimodal soft-intro VAE.

## B MOPOE - generalized ELBO

### Lemma 3

MoPoE generates a generalized ELBO with PoE and MoE being special cases.

To prove this lemma, first we need to show that ELBO_*MoPoE*_(*X*) is a valid multimodal ELBO, i.e. log *p*_*θ*_(*x*) ≥ ELBO_*MoPoE*_(*X*). Next we will show that MOPOE is a generalized case of both POE and MOE.

### Proof : MOPOE is a valid multimodal ELBO

In order to prove that ELBO_*MoPoE*_(*X*) is a valid multimodal ELBO, we need to show that log *p*_*θ*_(*X*) ≥ ELBO_*MoPoE*_(*X*). Note that while taking all possible subsets *X*_*k*_ ∈ *P* (*X*), we denote each subset by *k* for better readability.

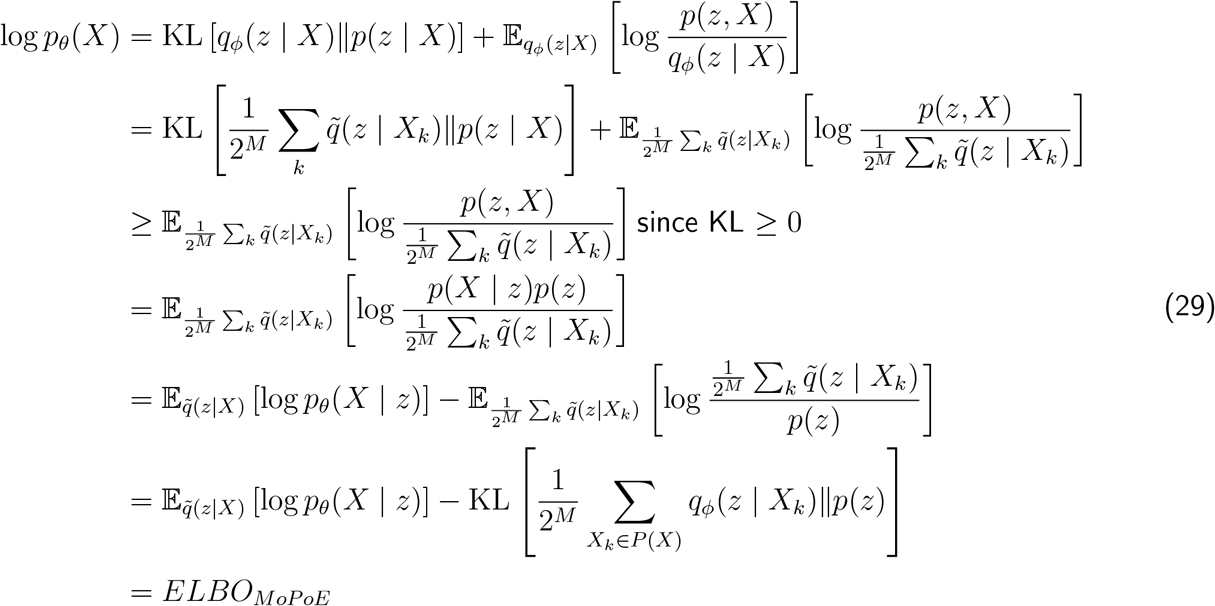

### Proof : MOPOE is a generalized version of POE and MOE

MoPoE generates a generalized ELBO with PoE and MoE being special cases. The MVAE architecture proposed by Wu and Goodman, 2018 only takes into account the full subset, i.e., the PoE of all data types. Trivially, this is a MoE with only a single component as follows:

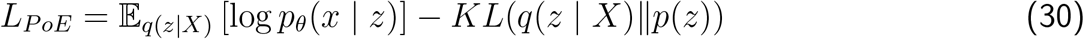

with

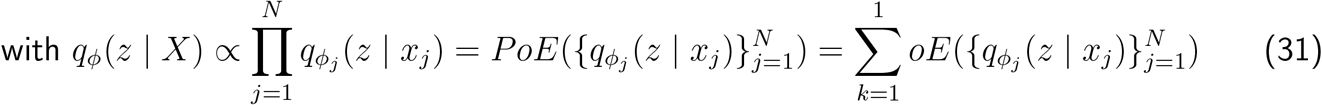

As the PoE of a single expert is just the expert itself, the MMVAE model (Shi et al., 2019) is the special case of MoPoE which takes only into account the N unimodal subsets as follows:

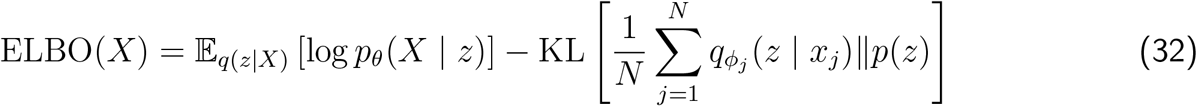

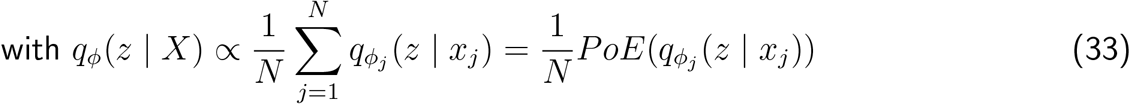

*L*_*MoE*_ is equivalent to a *L*_*MoPoE*_ of the N unimodal posterior approximations 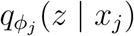 for *j* = 1, …, *N* . Therefore, the proposed MoPoE-VAE is a generalized formulation of the MVAE and MMVAE, which accounts for all subsets of modalities. The identified special cases offer a new perspective on the strengths and weaknesses of prior work: previous models focus on a specific subset of posteriors, which might lead to a decreased performance on the remaining subsets. In particular, the MVAE should perform best when all modalities are present, whereas the MMVAE should be most suitable when only a single modality is observed.

## Notes

### Competing Interest Statement

The authors have declared no competing interest.

